# Automated detection of sow posture changes with millimeter-wave radars and deep learning

**DOI:** 10.1101/2022.04.13.488188

**Authors:** Alexandre Dore, Mathieu Lihoreau, Jean Bailly, Yvon Billon, Jean-François Bompa, Edmond Ricard, Dominique Henry, Laurianne Canario, Hervé Aubert

## Abstract

Automated behavioural monitoring is increasingly required for animal welfare and precision agriculture. In pig farming, detailed analyses of sow activity are essential to identify and reduce the risks of piglets being crushed during postural changes of their mothers. Here we introduce a new, non-invasive, fast and accurate method for monitoring sow behaviour based on millimeter-wave radars and deep learning analysis. We used our method to predict postural changes in crated sows and distinguish the dangerous one that lie down abruptly from those that lie down carefully using transient postures. Two radars were placed on a metal backing above the head and the upper part of the back of each of ten sows to monitor their activity during 5 hours. We analysed the radar data with a convolutional neural network and identified five postures. The average sensitivity was 96.9% for standing, 90.8% for lying, 91.4% for nursing, 87.6% for sitting, but only 11.9% for kneeling. However, the average specificity and accuracy were greater than 92% for the five postures. Interestingly, two of the ten sows occasionally moved directly from standing to lying, without using the transient postures sitting and kneeling, thereby displaying risky behaviours for their piglets. Our radar-based classifier is more accurate, faster and require less memory than current computer vision approaches. Using more sows will improve the algorithm performance and facilitate future applications for large scale deployment in animal farming.

**Highlights:**

- Automated behavioural analysis is a major challenge for precision farming.
- We developed automated detection of lactating sow postures with radars and deep learning.
- We identified five postures, including transitions risky for the piglets.
- Our method is accurate, fast and requires less memory than computer vision.
- Radars thus hold considerable promises for high through-put recording of livestock activity.

## Introduction

An animal’s behaviour can inform about its stress level (von Borel et al. 2007) and personality (O’Malley et al., 2019). In the case of livestock, detailed behavioural are increasingly used to address key economic, ethical and societal issues related to animal health and welfare. In pig farming, for instance, sows are often confined in crates during the lactation period in order to limit their movements and slow down postural changes that can be dangerous for the piglets (Edwards and Fraser, 1997; Edwards et al. 2002). However, these rearing conditions are not ethically acceptable anymore. Long-term detailed behavioural monitoring of sow activity has thus become essential to move towards freer housing conditions during the lactation period (e.g. Chidgey et al., 2015; see Baxter et al. 2012 for a review). Ultimately, selecting sows that are calm, careful towards their piglets, and do not suffer from leg problems as future breeders would promote the use of more welfare-friendly systems.

The study of sow behaviour over extended periods of time is a fairly recent approach and technological solutions combining sensors and signal analyses for automated monitoring are being developed at a tremendous speed (Fernandes et al., 2020). Two broad categories of approaches have been used. Invasive technologies include sensors attached to the animals, such as accelerometers that record the acceleration and orientation of the sow across time. These sensors are typically embedded in an ear tag (Traulsen et al. 2018, Oczak et al 2020), a collar (Cornou et al., 2011), a harness (Canario et al., 2018), or fastened to one leg (Ringenbberg et al 2010). However, as piglets grow, they increasingly explore their environment and are likely to chew the device insistently, which may lead the sow to change posture and activity if the discomfort is significant.

Non-invasive technologies include remote sensors, such as infrared cells or video cameras. When positioned above a sow, infrared sensors enable the discrimination of lying, sitting and standing postures (Mainau et al. 2009). Cameras have been used to identify different postures when combined with machine or deep learning for data analysis (Zheng et al. 2018). Several image processing methods have proved effective in estimating the posture of sows kept in a crate (Bonneau et al. 2021, Nasirahmadi et al. 2019, Okinda et al. 2018). For instance, convolutional neural networks have been used to estimate the posture of a sow with its litter in an individual pen (Nasirahmadi et al. 2019). The rate of good classification for these methods is close to or above 90% for several classes of postures (e.g. sitting, standing, lying down). This suggest further development of algorithms for these non-invasive technologies can lead to the automatic monitoring of livestock behaviour at large scale in a near future. Pigs are good models to develop and crash test these approaches since sows are isolated from conspecifics during the lactation period. However, this seems out of reach with current computer vision approaches as the require large memories and computational power to be functional.

Here we introduce a non-invasive method for the automated detection of sow posture based on Frequency-Modulated Continuous-Wave (FMCW) millimeter-wave radars and illustrate its efficiency to identify postural changes of the sow that are risky for the piglets, when sows are maintained in crates. The ability of FMCW radars to monitor animal behaviour was recently demonstrated in insects (e.g. bees: Dore et al. 2020) and mammals (e.g. sheep: Henry et al. 2019; Dore et al. 2021). The main advantage of radars is that data collection and processing is fast and requires less memory than computer vision (the memory used for each measure is 1Mb and 5Gb for one image without compression). Manteuffel (2019) used Doppler radars to analyse sow activity, but limited their investigation to the detection of movements, since this kind of radars only gives the speed of the sow across time. In our study, we first compared the efficiency of the radar tracking system to state of art video tracking. We then applied a Convolutional Neural Network (CNN) to the data in order to classify the postures and measure the quality of posture prediction.

## Materials and Methods

### Sows and ground-truth measurements

Measurements were made on ten lactating sows from the Large White breed, on the GenESI INRAE experimental farm of Le Magneraud (France; doi: 10.15454/1.5572415481185847E12). Sows were studied up to two weeks after farrowing. Their behaviour was monitored with FMCW radars for a minimum of five hours. Video recordings were made in parallel to assess the reliability of the radar measurements. Postures were annotated manually from these recordings frame by frame every 100ms. The synchronisation between the radar and the video measurements was performed *a posteriori* by detecting manually the kneeling posture. This posture was chosen as reference because it was easily distinguished on the radar signal and was observed on all sows. We used five hours of annotated data per sow and considered five postures (Figure 1): standing; sitting; lying (i.e. ventrally); nursing (i.e. lying laterally with udder exposed), and kneeling.

**Figure 1.**
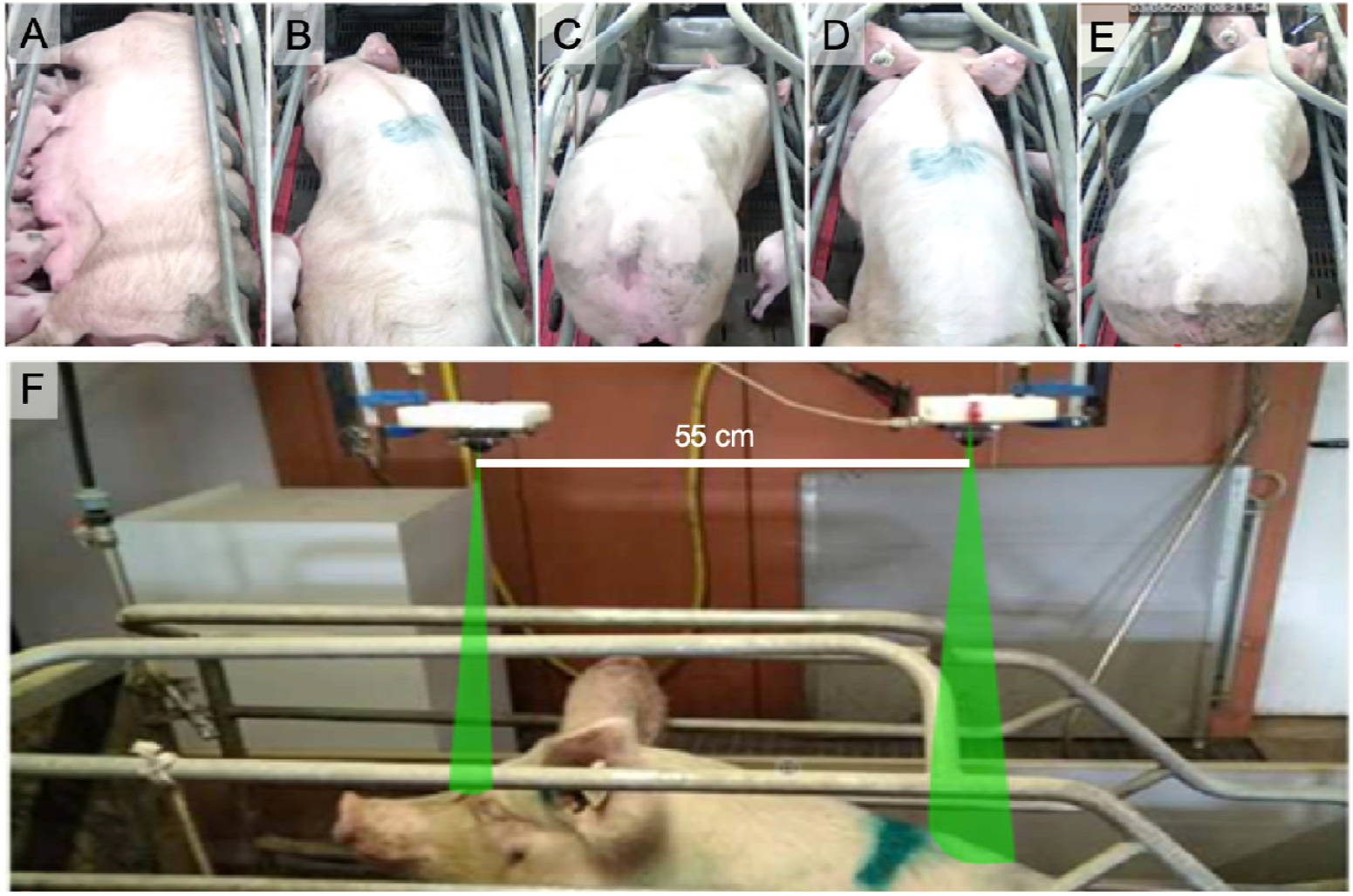
Images of sow postures as recorded by the video camera: A) nursing (lying laterally with udder exposed), B) lying (ventrally), C) kneeling, D) sitting, E) standing. F) Two FMCW radars were positioned above the sow to monitor head and back elevation. Radar beams are highlighted in green.

### Experimental Setup

Two millimeter-wave FMCW radars (EasyRadar) manufactured by Silicon Radar (Franckfurt, Germany) were used. The radars were attached on a metal structure in the middle axis of the crate (Figure 1). One radar was placed above the sow’s head (i.e.1.40m above ground). The other radar was located at the same height in the upper part of the sow’s back, at a distance of 55cm from the first radar. Each radar transmitted a frequency-modulated electromagnetic signal (*chirp*) every 5ms with the carrier frequency of 122GHz, a modulation bandwidth B of 6.9GHz, and a transmitted power of 4mW. The radars were used as altimeters with the theoretical depth resolution d=c/(2B), where c=3e8m/s is the speed of light in vacuum. A resolution of 2.1cm was sufficient to estimate the posture of the sow. Radars were small enough (100 × 70mm^2^) to be easily mounted above the crate without affecting the behaviour of the sows. A dielectric lens was mounted in front of each radar antenna to obtain a half-power beam width of 8°. In this way, the radars illuminated the sow, but not the top metallic bars of the crate.

### Radar signal analysis and electromagnetic clutter mitigation

The raw data (radar signal) are available upon request to the corresponding author. To estimate the posture of the sows, we analysed the signal delivered by the two radars over time. We computed the *beat frequency spectrum* (i.e., echo level magnitude as a function of the distance) by applying a Fourier transform (Stove et al. 1992) on the time signal. The result was a 2D raw data image of echo levels from which the posture of the sow could be estimated (Figure 2A). However, potential electromagnetic clutter, for instance due to the electromagnetic backscattering from the crate or from the piglets close to the sow, may alter the estimation accuracy of the posture. In order to mitigate this clutter, we used time-gating and spatial filtering. First, we applied a mean filter on the data across the slow time axis (Figure 2B). The mean was estimated over 1s, which corresponds to 20 consecutive radar measurements. Next, we applied Gabor filters to reduce the impact of uncorrelated data over time (Feichtinger and Strohmer 2012; see Figure 2C). Finally, we used an average filter (Figure 2D). The main advantage of using these filters is that they are simple to implement and suitable to real-time applications. In the final filtered 2D radar image (Figure 2D), some distinguishable behaviours of the sow could be detected at the distance of 1m, such as the transition to the standing posture around 1300s, 2700s and 3400s. These images obtained from two radars were used as input data for the posture classifier (see below).

**Figure 2.**
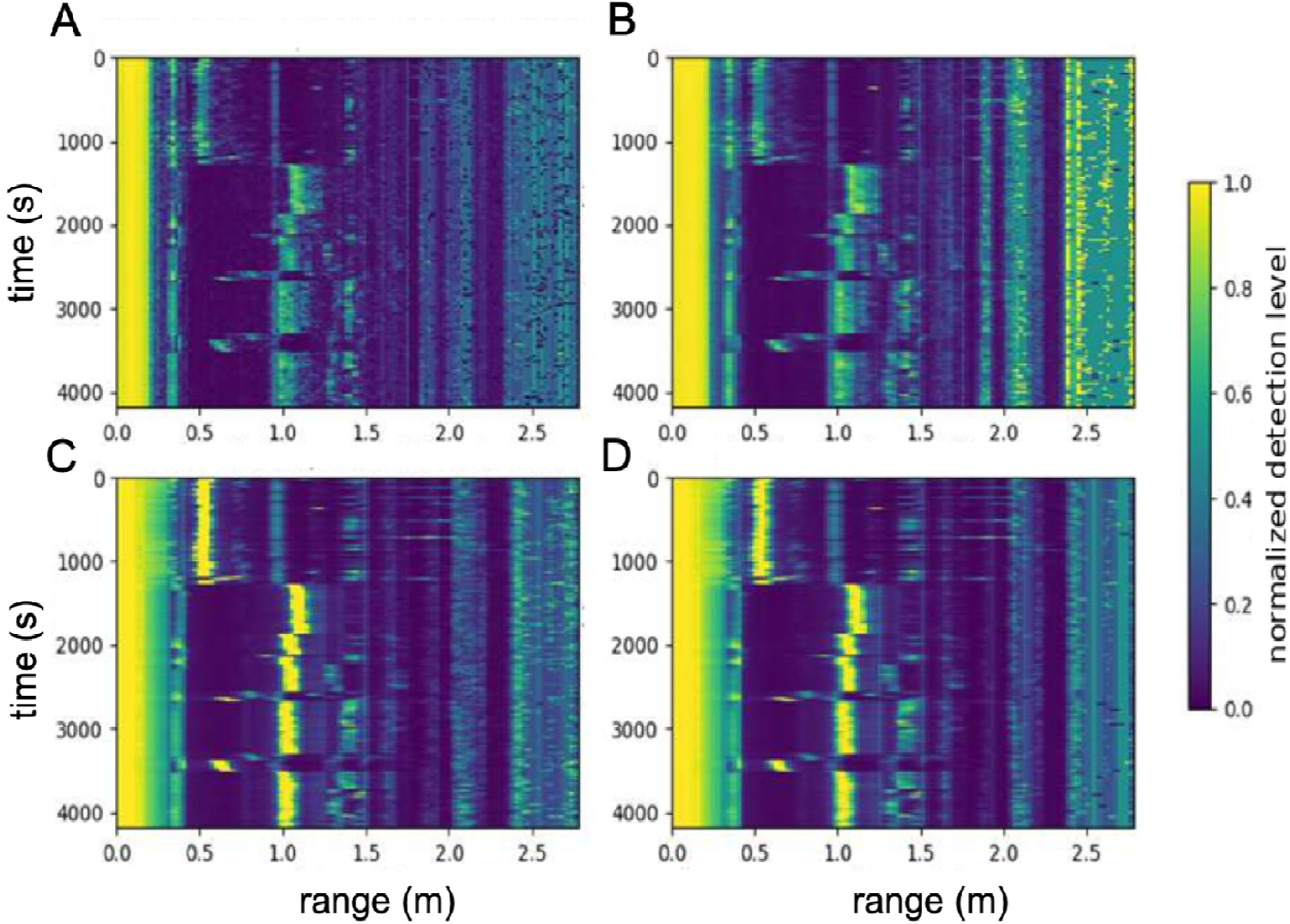
Outputs from the successive filters applied to the radar signal. A) Raw data; B) Temporal average filter; C) Gabor filter; D) Gabor filter plus average filter. Blue to yellow colours show low to high echo levels.

### Classification of sow postures using a Convolutional Neural Network analysis

To extract the postures of the sows, we used a CNN-based classifier. This approach has previously proved efficient for optical image classification (Simonyan and Zisserman 2015). Here the number of convolution layers as well as the number of filters per layer were chosen to speed up the training step of the model. A total of 16 filters were used for the first convolutional layers and the number of filters was doubled after each reduction phase (as proposed in Simonyan and Zisserman 2015). An illustration of the different layers of the network, including phases of convolution, activation, batch normalisation, pooling, dense, and dropout, is given in Figure 3.

**Figure 3.**
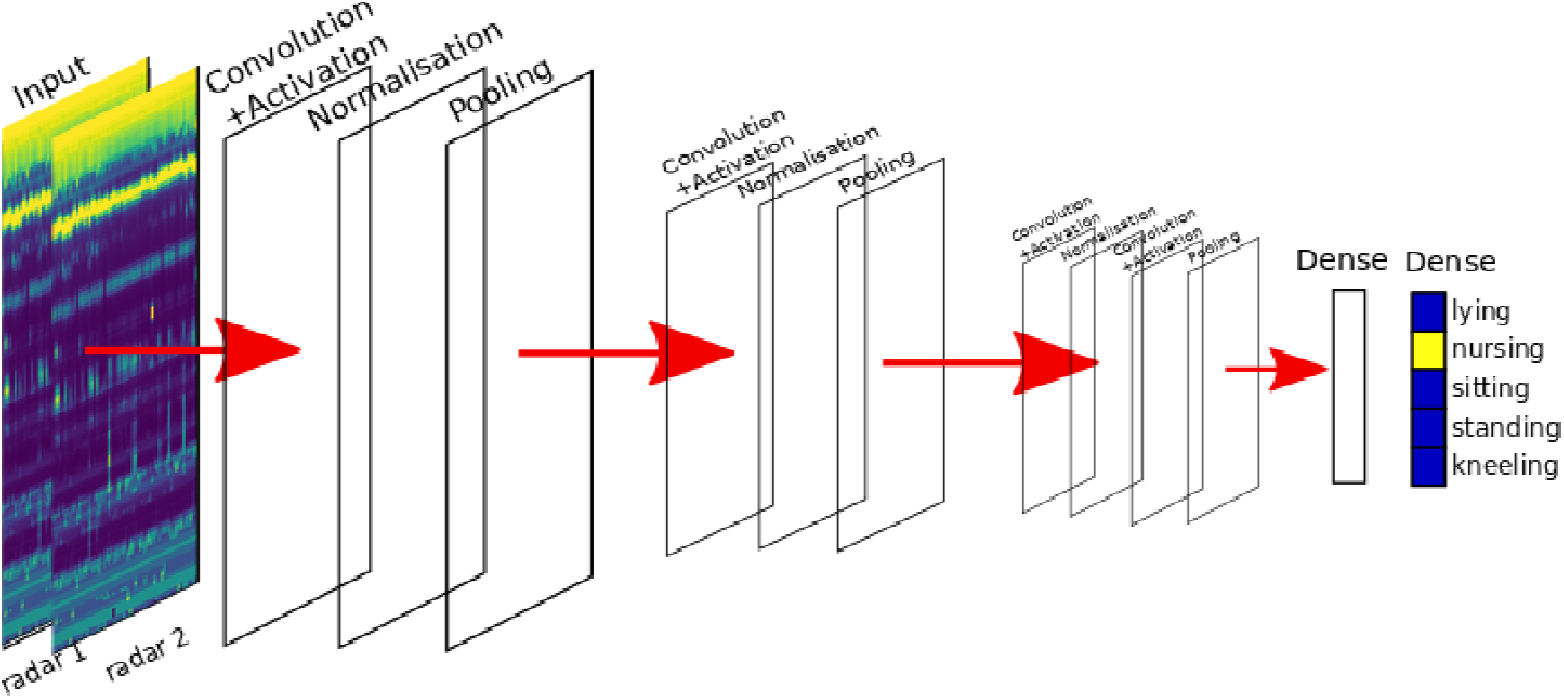
Illustration of the CNN model applied to radar images for estimating sow postures. Each convolution layer consists of a convolution filter of size 3×3. The pooling layer reduces the size of the data and dense layers are fully connected neurons.

CNN models were applied individually on each sow, since radar to ground distances were adjusted to the body size of each sow. To build a model, data from each sow was separated into two subsets containing respectively the first 60% and the last 40% of each posture data. The first subset was used for the training of the CNN. The second subset was used to estimate the quality of the predictions by checking that the results on the training dataset were not due to overfitting. During the training phase, we used 32 annotated data between two adaptations of the neural network weights. The CNN weights were adjusted following the method described by (Kingma and Ba 2014) which used a stochastic gradient descent and momentum estimation to optimise the neural network weights. The cost function was the standard function used by Murphy (2012) for multi-class classification. This function computes the minimal cross entropy for each output class. During the initial training, the five postures of interest were considered (Figure 1). After the training phase, each CNN was tested on the validation dataset.

The performance of the classifier in terms of the number of sow posture images that were correctly classified as well as the misclassified cases for each class was assessed with the confusion matrices. In the multi-class classification task, the evaluation metrics are computed based on these confusion matrices. We used three evaluation metrics to estimate the quality of the classifier: the specificity, the sensitivity and the accuracy (Fawcett 2006). The specificity (*sp*) indicates if the classifier gives negative results when the considered posture is not seen. The sensitivity (*sb*) indicates if the classifier gives positive results when the considered posture is seen. The accuracy (*acc*) indicates how often the classifier gives the true posture. Specificity, sensitivity and accuracy were calculated as follows:

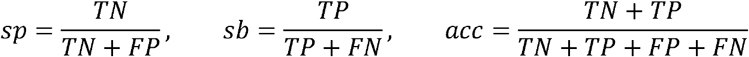

Where *TP* is the number of times that the estimated class corresponds to the true class, *TN* is the number of times the class is not predicted and corresponds to another class, *FP* is the number of times that the class is predicted, but does not correspond to the true class, and *FN* is the number of times that the class is not predicted and does correspond to the considered class.

## Results

For a detailed analysis of the quantitative results, the ten individual classification architectures were compared in the form of confusion matrices (global matrix for the ten sows, and individual matrices for each sow) shown in Figure 4. All occurrences of true/predicted postures are reported in the matrices for the 5 detected postures. The probability of false detection for each posture is derived from these first results. The specificity and sensitivity values of postures computed from the confusion matrices are reported in Table 1. Highest specificity values were obtained for sitting (99.3%) and standing (99.8%). On average, all postures had an accuracy and a specificity higher than 92%. Sensitivity was higher than 90% for all postures except kneeling that was detected in less than 10% of the time. Nursing and lying were confused by 10% on average. Kneeling had the lowest sensitivity (9.1%), but a high specificity (99.5%) and a high accuracy (99.4%). Our method can thus detect four out of five postures with a high specificity, a high sensibility and a high accuracy.

**Figure 4.**
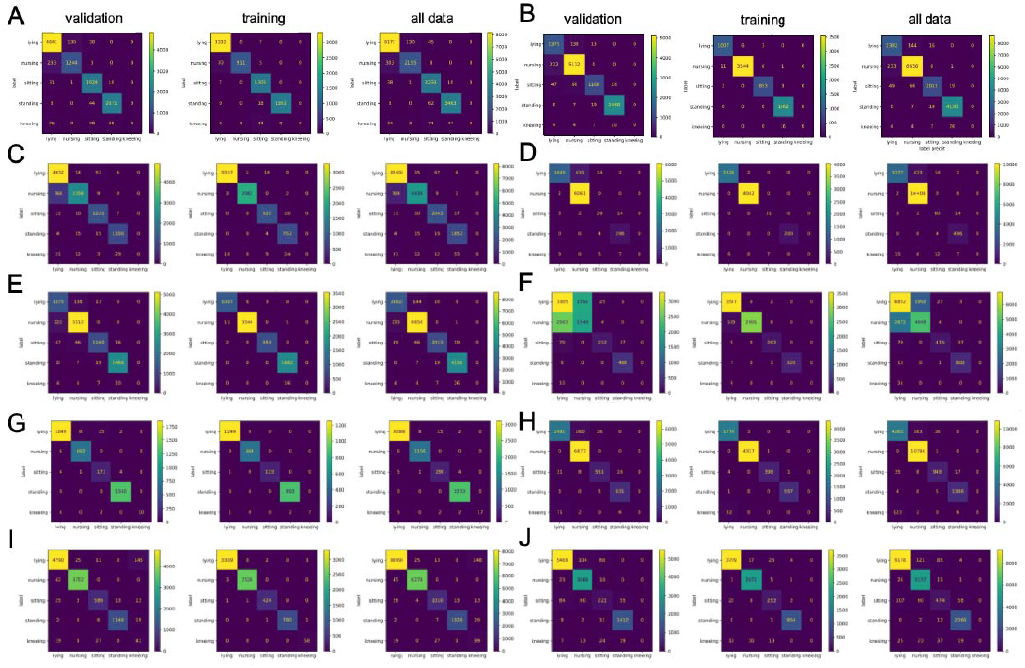
Confusion matrices for the 10 sows (A-J) and postures obtained from the individual CNN analyses. Vertical and horizontal axes indicate ground-truth measurement of the sow’s posture and the estimated posture, respectively.

**Table 1.** Specificity, sensitivity and accuracy of each of the 5 postures for the 10 sows obtained from individual CNNs. Labels A-J correspond to the same sows as in Figure 7.

|  | sensitivity |  |  |  |  | Specificity |  |  |  |  |
| --- | --- | --- | --- | --- | --- | --- | --- | --- | --- | --- |
| posture | lying | nursing | sitting | standing | kneeling | lying | nursing | sitting | standing | kneeling |
| sow A | 96.7 | 84.1 | 97.9 | 97.9 | 0.0 | 94.9 | 98.6 | 98.7 | 99.4 | 100.0 |
| sow B | 95.0 | 100.0 | 95.2 | 98.6 | 0.0 | 99.6 | 97.4 | 99.8 | 99.5 | 100.0 |
| sow C | 98.6 | 75.3 | 97.8 | 97.0 | 0.0 | 85.6 | 99.3 | 99.2 | 99.6 | 100.0 |
| sow D | 80.5 | 100.0 | 60.4 | 98.7 | 0.0 | 99.8 | 83.8 | 99.7 | 99.7 | 100.0 |
| sow E | 90.1 | 95.8 | 90.0 | 99.0 | 0.0 | 97.0 | 96.0 | 99.6 | 99.7 | 100.0 |
| sow F | 62.1 | 43.2 | 66.5 | 98.8 | 0.0 | 50.3 | 67.6 | 99.7 | 99.6 | 100.0 |
| sow G | 98.7 | 99.1 | 95.0 | 100.0 | 62.5 | 99.4 | 99.7 | 99.6 | 99.8 | 100.0 |
| sow H | 93.4 | 100.0 | 90.9 | 99.1 | 0.0 | 98.7 | 95.9 | 99.8 | 99.8 | 100.0 |
| sow I | 96.4 | 98.9 | 91.6 | 97.7 | 47.1 | 98.5 | 99.6 | 99.6 | 99.9 | 98.3 |
| sow J | 97.1 | 98.9 | 52.6 | 98.5 | 0.0 | 97.6 | 97.6 | 99.0 | 99.2 | 100.0 |
| Mean | 90.3 | 90.2 | 91.2 | 98.5 | 9.9 | 92.9 | 94.3 | 99.5 | 99.6 | 99.8 |
| Std | 10.9 | 17.4 | 16.3 | 0.8 | 22.2 | 14.5 | 9.8 | 0.4 | 0.2 | 0.5 |

The pie charts of the ground-truth measurements and estimated measurements of sow postures obtained from the validation datasets are shown in Figure 5. The relative time spent by each sow in each posture over a 2-hours period (corresponding to the validation dataset) was correctly estimated, with an error of less than 10% for all postures except kneeling. The difficulty in correctly estimating kneeling is likely due to the lack of sufficient radar data to analyse this posture with a convolutional neural network. Kneeling lasted only a few seconds (mean: 16s, standard deviation: 7s), which is considerably less than the other postures that lasted several minutes or hours (Table 2). In addition, the kneeling posture was much more sensitive than other postures to the measurement errors induced by the observer’s annotation delay (i.e. time needed to press a computer key to record the change in posture). Graphs of the probability of posture transitions indicate that the different sows lie down differently (Figure 7). In particular, two of them occasionally moved directly from standing to lying, without using the transient postures sitting and kneeling.

**Figure 5.**
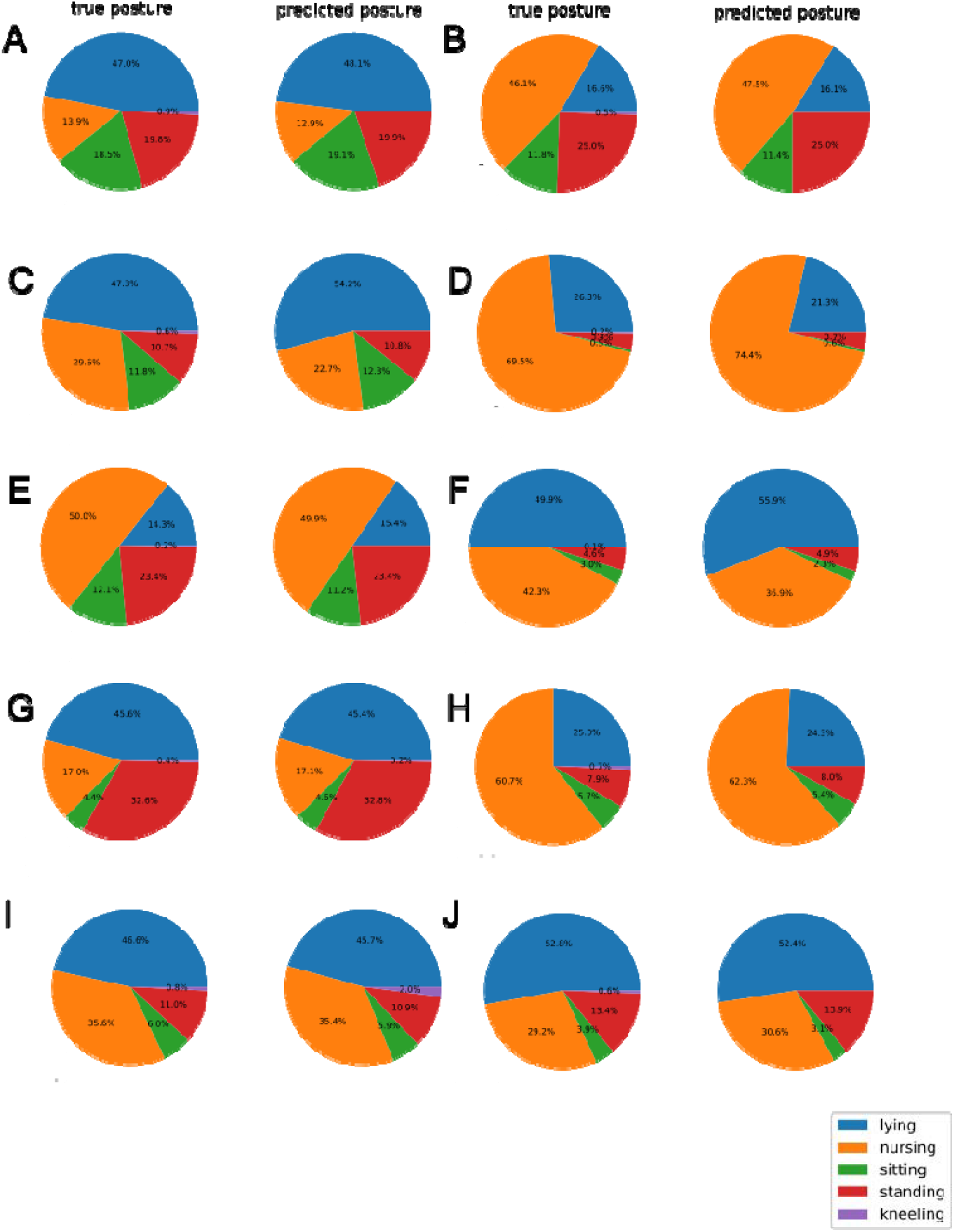
Percentage of time spent in the different postures by each sow (A-J) according to truth-ground measurement and estimated postures in the validation datasets. The higher th similarity between the true pie chart and the estimated pie chart of a sow, the better the prediction for that sow.

**Table 2.** Tme spent in each of the 5 postures by the 10 sows over the 5 hours period.

| Posture | total time spent (h) | rate of time spent (%) | mean of duration (min) | std of duration (min) |
| --- | --- | --- | --- | --- |
| lying | 16.66 | 33.56 | 7.19 | 5.99 |
| nursing | 22.67 | 45.67 | 20.31 | 30.51 |
| sitting | 3.76 | 7.57 | 1.72 | 2.23 |
| standing | 6.32 | 12.72 | 7.02 | 10.36 |
| kneeling | 0.24 | 0.48 | 0.29 | 0.14 |

## Discussion

Automated monitoring of behaviour in farm animals is a major challenge to improve animal welfare and breeding programs for genetic selection. In the case of pig farming, sow postural activity can be used as an indicator of health problems linked to increased piglet mortality. Here we introduced a new method for automated monitoring of postures and posture changes of sows kept in crate using the FMCW radars and a CNN classification. This solution is accurate, faster and requires little memory to store the data. It is thus applicable in real time. It also provides the advantages of not being affected by lightening condition (i.e. day and night analysis is never altered by the light at the level of the crate where the sow is measured) and non-invasive (i.e. do not alter sow behaviour in any way).

### Comparison with existing methods

Our radar-based tracking system provides several advantages over other existing methods. For instance, the radars are more accurate and less invasive than accelerometers (Ringgenberg et al. 2010; Canario et al., 2018). Although the detection sensitivity with the two approaches were similar for the three postures lying, nursing and standing, the radar data gave an estimation of the sitting posture with a much higher sensitivity than accelerometers (90% with radars, 37% with accelerometers). In addition, detecting other postures such as the sitting is meaningful for the analysis of sow behaviour related to piglet crushing, because it is included in the calculation of sow postural changes that are risky for the piglets (Melisová et al., 2014). Presumably, lying down through the transient postures sitting and kneeling indicates that sows are cautious towards their piglets when lying down.

We also found that the radar data processing is much faster and requires less memory than video processing for comparable results (Leonard et al., 2019; Bonneau et al. 2021). For example, Zheng et al. (2018) obtained similar results on the same five postures with a Kinect camera and a deep learning analysis on sows kept in individual pen, i.e., with more possibility to move. Using the same experimental design as ours, Bonneau et al (2021) achieved 98% sensitivity for four postures with a conventional 2D camera. Therefore, the main advantage of our approach lies in the processing of radar signal which is faster and requires less memory than that of video signal. Indeed, the processing by successive filtering is fast enough for a real-time application and this procedure is easily parallelisable. Using an Intel Core i7 processor with 4 threads makes it possible to process 12000 radar measurements per second. For a single measurement with two radars, only 800 bits are required, compared with several millions (3Mbits) for a standard definition camera. In our study the total data size of the five hours of measurements on ten sows presented was 2Go, which is compatible with the use of radars at large scale for farming applications. Additionally, by default and contrary to video image analysis, our analysis is independent of light conditions. This may be particularly relevant for future applications such as monitoring farrowing, which preferably takes place outside the working hours of the care takers, and therefore when the rooms are not lit (Canario et al., 2014).

### New possible analyses

Beyond being non-invasive, accurate and fast, the radar approach could be used to monitor sow carefulness when lying down (Pokorná et al., 2008). In the past, posture changes were rarely characterised by considering the passage through transient postures, i.e., kneeling and sitting (but see Pokorná et al., 2008; Wischner et al., 2010) and if so, have involved small groups of animals. Automated monitoring with radars in a farrowing crate enables such accurate recording of whether the sow lies down carefully. This feature can be linked to piglet crushing events.

Our data also indicate it is possible to detect piglets with radars. Prior to this study, preliminary measurements were performed in which the sow’s postures were estimated and the standard deviation was calculated over 20min, while the sow remained in a given posture. The standard deviation obtained was of 2.5cm for the nursing posture which is slightly higher than the theoretical depth resolution. However, the electromagnetic clutter may alter the measurement accuracy in some situations. For example, part of the metal bars of the crate that are visible to the radar may generate undesirable electromagnetic backscattering and result in false detections in the CNN analyses. This negative effect can be limited by precise positioning of the radars, so that the radar main beam does not intercept the metallic bars. False detections may also occur when piglets are close to the sow, or when they ride on its back. Figure 6 illustrates this phenomenon with two 2D radar images recorded simultaneously above (a) the head and (b) the back of a sow. The posture estimated from the radar processing (indicated as red cross) sometimes varies abruptly, due to the presence of piglets in the area of interest, i.e., on the back of the sow (e.g. around 600s, 800s and 1100s). Detection of the piglets at the udder is therefore presumably possible. The radars can be effective in monitoring additional behavioural aspects of the sow-piglets relationship that are of importance for piglet survival.

**Figure 6.**
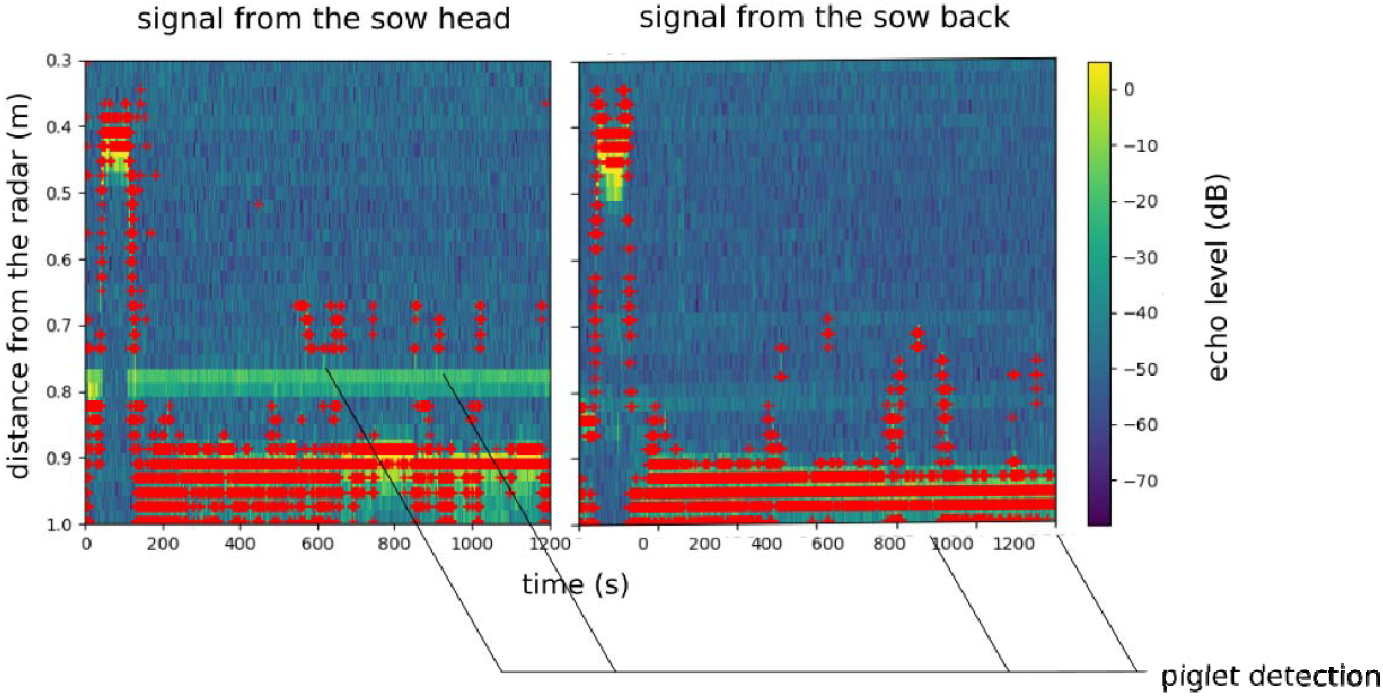
Two-dimensional images of the two radars illuminating (a) the head and (b) the back of a sow when piglets are riding on its back. Red crosses represent the computed first local maxima for each slow time bin. The presence of piglets on the sow back can be observed at various times. A threshold for the distance estimation can be used (around 0.6m for the two radars) to determine if the sow is lying or standing.

**Figure 7.**
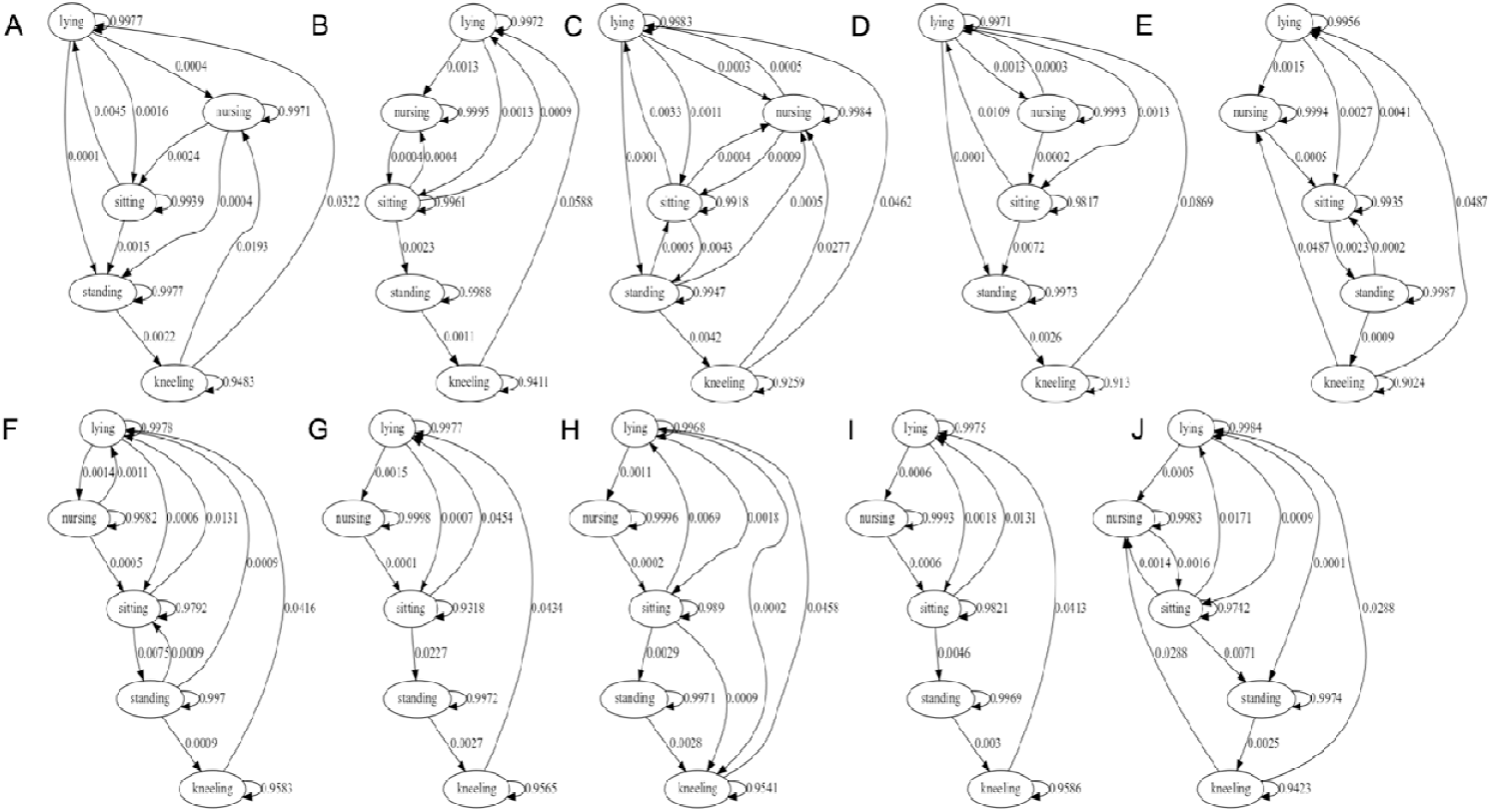
Graphs of the probability of posture transitions, i.e., the probability of switching between the 5 postures for the 10 sows (A-J) over a 5 hours period. Labels A-J correspond to the same sows as in Table 1.

### Conclusions and perspectives

Our study demonstrates the applicability and efficiency of FMCW radars to monitor postures and postural changes of crated sows. This approach holds considerable promises for future applications since minor developments are possible to improve the system. For instance, adding radars will enable to detect new postures and consolidate the estimation of the current ones. Our method was applied to detect the postural activity of the sow from the estimation of the height of its head and its back. Consequently, it could not detect lateral movements of the sow and if the sow was lying on the right or left side. This estimation would be possible by adding a third radar on the side of the sow for detecting lateral movements and consequently, for estimating the orientation of the sow in the crate and whether she is nursing the piglets or not. Moreover, the kneeling posture was not correctly estimated for any sow. This posture is interesting to capture, as it reflects a cautious approach of the sow to lie down, compared with sows that slide along the bars from the standing posture (Pokorná et al., 2008). Sliding is a less suitable way to lie down for sows as it induces some piglet crushing (Fraser 1990, Damm et al. 2005). The kneeling posture lasts only a few seconds, and its estimation is sensitive to potential annotation errors due to human delay between observation and pressing a computer key. However, from the monitoring of more sows and fixing the radar on the crate rather than a metallic structure above, it should be possible to improve the reliability of the measurement and the accuracy of the posture estimator.

Using different types of radars can also improve piglet detection. With our method, the detection of piglets on the floor is difficult because they rarely enter the radars field of view and are small compared to the sow. The radar beam width was too small to allow detecting piglets. One solution to this problem would be to use two 2D radars instead of the current device, and position one on each side of the sow. In this way, the detection should work even if the sow hides the piglets from one of the radars beam. Access to such data on the presence of piglets around the sow would benefit the characterisation of sow carefulness to lie down (Pokorná et al., 2008; Wischner et al., 2010). Adjusting radar characteristics can also increase detection distances. Until now, results were obtained in indoor environment where the sow cannot move. The radar detects the back of the sow at all times. The use of radar with sows free of movements can be more challenging. Then, it should be possible to detect the sow by using a single 3D radar with a larger beam width, including position and posture simultaneously. To do so, the radar should be placed centrally on the ceiling above the pen to cover the entire pen with the radar beam.

Finally, adding data would also make it possible to define a single classifier to estimate the posture for all sows and develop more detailed behavioural analyses (Bonneau et al., 2021). So far, the relevance of radars to detect sows that are injured (e.g. excessive time spent lying) or that are not careful towards their piglets (e.g. fast transition from standing to lying) has not been studied. Data recorded for longer periods on animals contrasted on these aspects could be collected with the radar technology. All these improvements are easily reachable, which strongly suggests that automated fine recording with radars will help to analyse the relationship between sow postural changes and risk of piglet crushing. Importantly, radar tracking has limited constraints on the size and shape of animals (Henry et al. 2018; Dore et al. 2020; Dore et al. 2021), thus offering ample possibilities for tracking a wide range of animals in various environments in applied and fundamental ethology research.

## Acknowledgements

We thank the staff of the GenESI INRAE experimental farm for their practical help and care of the animals.

